# Mitochondrial RNA processing promotes translation by resolving structured precursor RNAs

**DOI:** 10.64898/2026.07.30.741833

**Authors:** Claire H. Nuessmeier, Gyan Prakash, Lisa N. Hansen, Mary T. Couvillion, L. Stirling Churchman

## Abstract

Mammalian mitochondrial mRNAs (mt-mRNAs) are excised from polycistronic precursors primarily through cleavage of flanking tRNAs. A subset of junctions lacks intervening tRNAs and is processed by a non-canonical mechanism involving members of the FASTK family of RNA-binding proteins; however, the consequences of non-canonical processing for downstream gene expression remain poorly understood. Long-read RNA sequencing of FASTKD4-deficient cells revealed selective processing defects at the *ATP8/6-CO3* and *ND5-CYB* junctions. Although processing of both junctions was impaired, mitoribosome profiling showed that only *CYB* translational efficiency declined, explaining reduced CYB protein despite unchanged *CYB* mRNA abundance. Mitoribosomes preferentially initiated on processed *CYB* transcripts despite the accumulation of unprocessed precursors, but engaged both processed and unprocessed *CO3* transcripts. Chemical probing of RNA structure revealed that the unprocessed *ND5-CYB* junction contains a predicted RNA secondary structure adjacent to the *CYB* start codon, while the *ATP8/6-CO3* junction is unstructured. Our results show that FASTKD4-dependent processing of the *ND5-CYB* precursor is required for efficient *CYB* translation and support a model in which non-canonical processing promotes translation of select transcripts by removing precursor RNA structures that hinder mitoribosome engagement.

## Introduction

The mammalian mitochondrial DNA (mtDNA) encodes thirteen essential subunits of the oxidative phosphorylation (OXPHOS) complexes, as well as the rRNAs and tRNAs required for their expression, on a compact, 16.5 kilobase genome (1). When transcribed, the mtDNA yields two long, polycistronic transcripts which must be efficiently processed into mature rRNAs, tRNAs, and mRNAs (Fig. 1A). Impaired processing of the polycistrons is deleterious for respiratory function, establishing mitochondrial RNA (mtRNA) processing as a vital step in OXPHOS gene expression (2–4).

**Figure 1:**
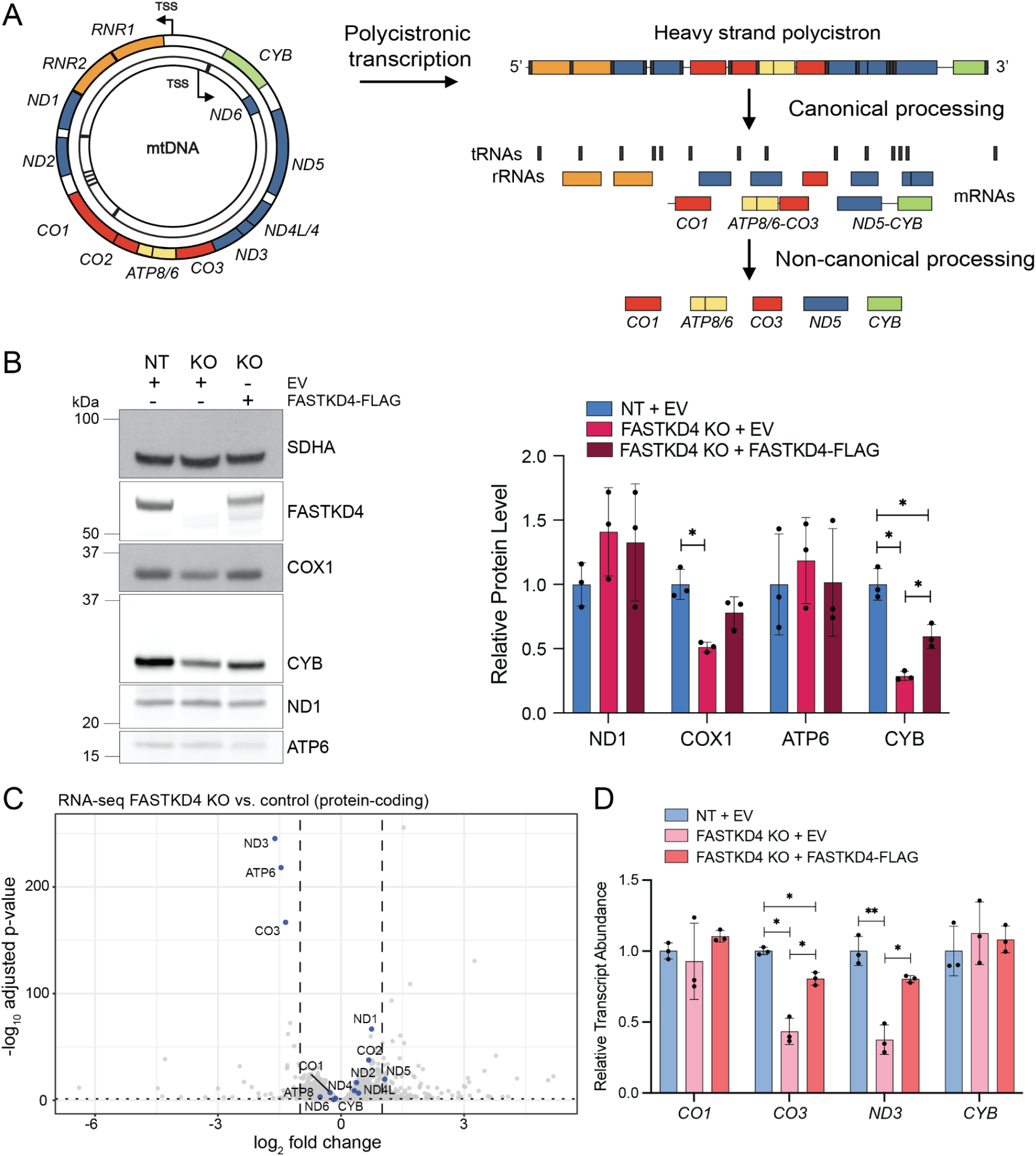
Loss of FASTKD4 differentially affects mtDNA-encoded gene products. **A)** Schematic of human mitochondrial DNA expression. mt-mRNAs are colored by OXPHOS complex. mt-rRNAs are shown in orange and mt-tRNAs in dark grey. TSS, transcription start site. **B)** Representative Western blot analysis of OXPHOS subunits (left) and quantification of protein levels (right) from three replicate measurements of whole cell lysate from NT, FASTKD4 KO, and FASTKD4 KO rescue cells. Error bars represent standard deviation. Brown-Forsythe and Welch ANOVA test followed by Dunnett’s T3 multiple comparisons test, and significant p-values are indicated; *p-value < 0.05. kDa, kilodalton; EV, empty vector. **C)** Volcano plot depicting log2 fold change (FASTKD4 KO/NT control; x-axis) and -log10 adjusted p-value (y-axis) from RNA-seq. mt-mRNAs are highlighted. **D)** NT-normalized transcript abundance (relative to ACTB) for a subset of mt-mRNAs as quantified by RT-qPCR for three replicate measurements from NT, FASTKD4 KO, and FASTKD4-FLAG rescue cells. Error bars represent standard deviation. Brown-Forsythe and Welch ANOVA test followed by Dunnett’s T3 multiple comparisons test, and significant p-values are indicated; *p-value < 0.05 and **p-value < 0.01.

The mitochondrial rRNAs and a majority of the mRNAs are flanked by tRNAs (5). Recognition and cleavage of the tRNAs result in the liberation of the mature rRNAs and mRNAs from the polycistron (6). As most mRNAs are flanked by tRNAs, this ‘tRNA punctuation’ model of processing is the canonical processing mechanism. The nuclear-encoded RNase P and RNase Z complexes cleave the 5′ and 3′ ends of the tRNAs, respectively (7–10). Processing by these complexes generates mature, leaderless mRNAs with minimal or absent 5′ untranslated regions (UTRs).

A subset of junctions, including the 5′ *CO1, ATP8/6-CO3,* and *ND5-CYB* junctions, lack an intervening tRNA, and their maturation therefore, requires an alternative, non-canonical processing mechanism (Fig. 1A). Cleavage at these junctions generates mature downstream mitochondrial mRNAs (mt-mRNAs) with start codons at or near their 5′ ends: *CYB* and *CO3* are leaderless, whereas *CO1* retains a three-nucleotide 5′ UTR. Two members of the Fas-activated serine/threonine kinase (FASTK) family of mitochondrial-localized RNA-binding proteins, FASTKD4 and FASTKD5, have been proposed to be involved in the processing of non-canonical junctions. Loss of FASTKD4 or FASTKD5 leads to increased abundance of unprocessed non-canonical transcripts; depletion of FASTKD4 primarily affects processing of the *ATP8/6-CO3* and *ND5-CYB* junctions, while loss of FASTKD5 impacts processing of all three non-canonical junctions and is especially deleterious for *CO1* maturation (11–16).

Similar to transcripts flanked by tRNAs, non-canonical transcripts exist predominantly in their processed forms in wild-type cells (13, 15). Unlike canonical cleavage, non-canonical processing is not required to generate the full complement of mitochondrial tRNAs. Therefore, the necessity for non-canonical processing and how retention of non-canonical junctions affects downstream gene expression remains unclear. Because mt-mRNAs are transcribed as a polycistronic precursor, most regulation of the mt-mRNA lifecycle is post-transcriptional: occurring at the levels of RNA turnover, mitoribosome engagement, or translation. Although defective non-canonical processing alters OXPHOS protein expression (13, 15), how this defect propagates through these post-transcriptional stages, and whether each junction is affected similarly, remains unclear.

We addressed these questions in FASTKD4-deficient human cells, tracking the lifecycle of mt-mRNAs from processing to translation. Using long-read RNA sequencing, we show that loss of FASTKD4 selectively impairs processing at the *ND5-CYB* and *ATP8/6-CO3* non-canonical junctions while canonical junction processing remains largely unchanged. Impaired processing coincides with reduced abundance of CYB protein. Despite processing defects at the *ND5-CYB* and *ATP8/6-CO3* junctions, mitoribosome profiling reveals that translational efficiency is reduced specifically for *CYB*, whereas *CO3* translational efficiency is maintained. Chemical probing of RNA structure indicates that the unprocessed *ND5-CYB* junction harbors a predicted secondary structure preceding the *CYB* start codon, while the *ATP8/6-CO3* junction remains unstructured. Together, our results reveal that non-canonical processing permits translation in a junction-specific manner and removes structural barriers within precursor transcripts.

## Methods

### Key reagents

Supplementary Table 1 contains a list of antibodies, qRT-PCR probe sets, oligonucleotides, and cell lines used in this work.

### Cell culture

HEK293T and HeLa cells were grown in high glucose Dulbecco’s modified Eagle medium (DMEM) supplemented with 10% standard fetal bovine serum (FBS; Thermo Fisher Scientific) and were cultured without penicillin or streptomycin. Medium for HEK293T LRPPRC KO cells was supplemented with sodium pyruvate (1 mM) and uridine (50 µg/mL) to maintain cells with dysfunctional mitochondria. Cells were maintained in a humidified incubator at 37°C with 5% CO_2_. Cells were regularly tested for mycoplasma contamination using a PCR-based mycoplasma detection kit (ATCC).

### Lentivirus packaging, transductions, and CRISPR KO

Lentiviruses were packaged in wildtype HEK293T cells following standard protocol for third generation packaging. Lentivirus-containing media was collected 48 hours following transfection of HEK293T cells and filtered using 0.45 μm PES membranes or centrifuged for 5 minutes at 300*g* to remove cellular debris. HEK293T cells were transduced by direct addition of lentiviral supernatants to cell cultures (1:1 ratio of viral supernatant to cell culture medium) in the presence of 8 µg/mL polybrene for 16–24 hr then replaced with fresh media.

CRISPR-mediated knock-outs of target genes was performed by transducing HEK293T and HeLa cells with lentivirus expressing Cas9 and lentivirus expressing sgRNAs targeting genes of interest. sgRNAs targeting the adeno-associated virus site 1 (AAVS1) locus served as a non-targeting (NT) control. Sequences of sgRNAs can be found in Supp. Table 1. Lentiviral transductions were performed as described above, followed by selection with 2 µg/mL puromycin for 72 hr. For HEK293T, single-cell clones were obtained by dilution of pooled mutant cells to one cell per well in 96-well plates, clonal expansion for 2-3 weeks, and confirmation of homozygous disruption of genes of interest by PCR and Western blotting.

### Cell lysis and RNA extraction

Cells were washed with ice-cold PBS before lysis or RNA extraction. Cell lysates were prepared in radioimmunoprecipitation assay (RIPA) buffer in the presence of protease inhibitors, unless otherwise specified. Lysates were cleared by centrifugation at 21, 000*g* and RNA extraction was performed using Trizol and isopropanol precipitations, following manufacturer’s instructions. RNA was DNaseI-treated using amplification grade DNaseI (Invitrogen) by the following method: 5 µg of RNA was combined with 2 µL 10X DNaseI reaction buffer, 2 µL DNaseI, and nuclease-free water up to 20 µL. The solution was incubated at 37°C for 15 min, 2 µL 25 mM EDTA was added, and the solution was heated at 65°C for 10 min.

### Immunoblotting and antibodies

Protein lysates for Western blotting were prepared in radio immunoprecipitation assay (RIPA) buffer in the presence of protease inhibitors and lysates were normalized by bicinchoninic acid (BCA) assay. Standard sodium dodecyl-sulfate (SDS)-PAGE and electroblotting protocols were used and samples were run on NuPAGE 4-12% Bis-Tris gels (Thermo Fisher). Samples were not boiled prior to SDS-PAGE when analysing mitochondrial-encoded proteins. This work used the antibodies found in Supp. Table 1 at the following dilutions in 5% skim milk in TBST MT-CO1/COX1 (1:2, 000), ACTB (1:5, 000), MRPS18B (1:1, 000), MRPL12 (Proteintech, 14795-1-AP, 1:1000), SDHA (1:15, 000), MT-ATP6 (1:1, 000), MT-ND1 (1:1, 000), MT-CYB (1:10, 000), MT-CO2/COX2 (1:2000), TBRG4 (1:2, 000).

### CellTiter-Glo

Relative ATP production by HEK293T NT control and FASTKD4 KO cells under glycolysis or ATP synthase inhibition (Oligomycin) was assayed by CellTiter-Glo 2.0 (Promega). Cells were cultured with 10 mM glucose, 10 mM 2-deoxy-D-glucose, or 100 nM oligomycin for 3 days before CellTiter-Glo assay was performed following manufacturer’s instructions.

### Annexin V Staining assay

HEK293T NT and FASTKD4 KO cells were seeded at 150K cells per well in 6-well plates. The following day, cells were treated with 10 mM 2-deoxyglucose or left untreated. Following 4 days of treatment, cells were harvested for Annexin V staining (Pacific Blue Annexin V/SYTOX AAdvanced Apoptosis Kit for Flow Cytometry, Invitrogen) and cell analysis by flow cytometry. Annexin V staining was performed following manufacturer’s instructions. Flow cytometry was performed with a BD LSR-II analyzer (Harvard Medical School Immunology Flow Core) and data analysis was performed with FlowJo.

### Mitochondrial isolation

HEK293T cells were grown to 80-90% confluency in 500 cm^2^ square TC-treated culture dishes (Corning). To isolate mitochondria, each plate was washed twice with cold PBS before being lysed with 3 mL of freshly prepared hypotonic mitochondrial isolation buffer (MIB; 50 mM HEPES-KOH pH 7.5, 10 mM KCl, 1.5 mM MgCl_2_, 1 mM EDTA, 1 mM EGTA, 1 mM DTT, 1x EDTA-free protease inhibitor cocktail; (17)) and left to swell on ice for 5 min. ⅓ volume of sucrose-mannitol buffer (SM4; 280 mM sucrose, 840 mM mannitol, 50 mM HEPES-KOH 10 mM KCl, 1.5 mM MgCl_2_, 1 mM EDTA, 1 mM EGTA, 1 mM DTT, 1x EDTA-free protease inhibitor cocktail; (17)) was added and cells were lysed with a 27 G 1 ½” needle. The lysate was spun at 800*g* for 10 min at 4°C to pellet unbroken cells. The resulting supernatant was spun at 10, 000*g* for 10 min to pellet the mitochondria. The mitochondrial pellet was washed once with MIBSM buffer (3:1 ratio of MIB:SM4 buffers; (17)) before further application.

### Nanopore long-read RNA sequencing

Total RNA was extracted as described previously. poly(A)+ mRNAs were enriched for using the Dynabeads mRNA Purification Kit (ThermoFischer, 61006), following manufacturer’s instructions. Long-read RNA sequencing library preparation was performed using the kit SQK-RNA002 (Oxford Nanopore Technologies) with 500 ng of RNA according to manufacturer’s instructions with the following exceptions: the RCS was omitted from the initial ligation mix and replaced with nuclease-free water and the ligation of the reverse transcription adapter (RTA) was performed for 15 min rather than 10 min. Libraries were sequenced on a MinION device (Oxford Nanopore Technologies) for up to 72 hours. Sequencing reads were basecalled in real time using MinKNOW (release 20.10.3 or later), and reads with a mean basecalling quality score greater than 7 (<20% base-calling error) were retained for downstream analysis.. Uracil bases in filtered reads were converted to thymine, and reads were aligned to the hg38 human reference genome using minimap2 with the parameters with parameters -ax splice -uf -k14 (18). The resulting SAM alignments were converted to coordinate-sorted BAM files and indexed using SAMtools. Multi-mapping reads were included in all downstream analyses. Mitochondrial read alignments were intersected with annotated mitochondrial transcripts in a strand-specific manner. Reads were required to overlap an annotated transcript by at least 25 nucleotides (nt) for transcript-assignment analyses. Transcript abundance was assessed using either the complete annotated transcript or windows encompassing the terminal 100 or 200 nt of each transcript. Duplicate read–transcript assignments were removed before counting.

Determination of processing was performed as in McShane et al. (19). Briefly, the 5′ and 3′ ends of each read mapping to mt-mRNAs were evaluated relative to the following windows spanning -15 to +50 nt from the transcript start or -15 to +15 nt from the transcript end, respectively. Reads beginning in the transcript start window were classified as ‘5′ processed’ and reads terminating in the transcript end window were classified ‘3′ processed’. Reads which covered the downstream gene body and which overlapped at least 15 nt with the window spanning -60 to -15 nt relative to the transcript start were classified as ‘5′ unprocessed’. Reads which covered the upstream gene body and which overlapped at least 15 nt with the window spanning +15 to +60 nt relative to the transcript end were classified as ‘3′ unprocessed’. Nanopolish was used to estimate poly(A)-tail lengths from the raw FAST5 signal using the corresponding basecalled FASTQ reads, coordinate-sorted BAM alignments, and reference genome. For mitochondrial transcript-level analyses, reads were additionally required to have a mapped 3′ end within ±15 nt of the annotated transcript end (20).

### RNA-sequencing

Total RNA was extracted as described previously and DNaseI-treated. 681 ng of DNase-treated RNA per sample was submitted to Plasmidsaurus for 3′ end strand-specific RNA-sequencing. Libraries were sequenced on an Illumina platform and processed using the Plasmidsaurus bioinformatic RNA-seq pipeline. In brief, raw base call files were converted to FASTQ and demultiplexed with BCL Convert (v4.3.6) and fqtk (v0.3.1). Reads were filtered using FastP (v0.24.0; poly-X tail trimming, 3′ end quality-based tail trimming, minimum Phred quality score of 15, minimum length of 50 bp). STAR aligner (v2.7.11) was used to align filtered reads to the reference genome (*Homo sapiens* GRCh38 Ensembl release 114) with removal of non-canonical splice junctions and output of unmapped reads. Coordinate sorting of BAM files was performed with samtools (v1.22.1). Reads were de-duplicated using UMICollapse (v1.1.0). RSeQC (v5.0.4) and Qualimap (v2.3) were used to evaluate alignment quality and a comprehensive QC report was created with MultiQC (v1.32). Libraries generated a median of 19 million filtered reads per sample (range from 17-20 million), calculated as the sum of uniquely mapped, multi-mapped, and unmapped reads. A median of 64.7% reads uniquely mapped to the reference genome. After unique molecular identifier (UMI)-based deduplication, a median of 9.8 million deduplicated reads per sample (range 9.3-10.7 million) remained for downstream analysis.

Downstream differential expression analysis was performed independently in R (v4.4.2) using DESeq2 (v1.50.0). Gene-level counts were first obtained with featureCounts (Subread v2.20.0) and the GENCODE v48 GRCh38.p14 GTF annotation. Reads overlapping annotated exons were summarized by gene_id in strand-specific mode. Chromosome naming in the annotation file was modified to match BAM headers. Genes with low counts were filtered prior to DESeq2 analysis: genes with at least 10 reads in at least 3 of the 9 samples (NT, KO, rescue in triplicate) were retained. Counts were normalized with the default DESeq2 median-of-ratios method. Gene-wise dispersions were estimated, and negative binomial generalized linear models were fitted. Pairwise contrasts were generated for FASTKD4 knockout (KO) versus non-targeting (NT) control, FASTKD4-FLAG rescue versus NT, and rescue versus KO. Statistical significance was determined by the Wald test and P-values were adjusted using the Benjamini-Hochberg correction. Genes with adjusted P-value (padj) < 0.05 were considered differentially expressed. For visualization, log2 fold-change estimates were shrunk with apeglm (v1.28.0) and gene annotations were included using biomaRt (v2.62.1; (21–23).

### RNA abundance and turnover measurements with MitoStrings

mtRNA turnover was determined by quantifying the decrease in old/unlabeled mtRNA following a 4-thiouridine (4sU) pulse, as described in McShane et al. and Ietswaart et al. (19, 24). HEK293T cells were labeled with 500 µM 4sU for 90 minutes, or left unlabeled, then lysed in Trizol and RNA was extracted as described previously. *In vitro* transcribed ERCC-0048 was used as a spike-in and was added during the RNA extraction. 20 µg RNA was denatured at 60°C for 10 min, and then rested on ice for 2 min prior to biotinylation. All work prior to biotinylation was done in the dark or with the lights turned off to protect the photoreactive 4sU. RNA was biotinylated by the addition of 16 µg biotin-MTS (Biotium) in 20% dimethylformamide (Sigma), 20 mM HEPES pH 7.4, and 1 mM EDTA and incubation for 30 min at 24°C with shaking at 800 rpm. Free biotin was removed by spinning samples with chloroform:isoamyl alcohol (24:1; Sigma) in phase-lock heavy gel tubes (5prime) following the manufacturer’s instructions and RNA was again precipitated. 2 µg of RNA was incubated with µMACS streptavidin beads from µMACS Streptavidin kit for 15 min at 24°C with shaking at 800 rpm to remove biotinylated RNA. RNA and beads were loaded onto a µMACS column, beads were washed with 100 mM Tris pH 7.5, 10 mM EDTA, 1 M NaCl, 0.1% Tween 20, and unlabeled RNA was collected in the flowthrough. Flowthrough RNA was purified using miRNeasy kit (Quiagen) following the manufacturer’s protocol, including DNaseI treatment. 50 ng of RNA was incubated at 67°C for 16 hrs with XT Tagset-24 (NanoString Technologies) and with DNA-probes specific for mitochondrial RNA (MitoString-probes modified as in Isaac et al. and McShane et al.) in hybridization buffer (NanoString Technologies) according to the manufacturer’s instructions (19, 25). Samples were then loaded onto nCounter Sprint Cartridge and quantified using the nCounter SPRINT Profiler (NanoString Technologies) at the Boston Children’s Hospital Molecular Genetics Core. Data was processed using nSolver and RStudio.

### Rescue experiments and transient transfections

1.5x10^5^ HEK293T NT and FASTKD4 KO cells were seeded in 6-well plates for transient transfection. One day after seeding, cells were transfected with 1 µg of either FLAG-tagged FASTKD4 that had been cloned into the pcDNA3.1 vector (Addgene) or empty vector. Transient transfection was carried out using Lipofectamine 3000 (Thermo Fischer) following manufacturer’s instructions. 2 µL of P3000 reagent was used per 1 µg of plasmid and 1.5X P3000 volume of Lipofectamine 3000 reagent was added. Cells were harvested 72 hrs after transfection for RNA extraction or cell lysis and immunoblotting, as described previously.

### RT-qPCR

For RT-qPCR quantification of mRNA levels, 1.82 µg of DNaseI-treated total RNA was reverse transcribed using a SuperScript III First-Strand Synthesis kit (Invitrogen). cDNA was diluted 1:100 and 2 µL was used for Taqman gene expression assays using TaqMan Fast Advanced Master Mix, no UNG (Thermo Fisher Scientific) and the probe sets found in Supp. Table 1. RT-qPCR was performed using a CFX Opus 384 Real-Time PCR System (BioRad) with the following conditions: 95°C for 20 sec, 40 cycles (95°C for 3 sec, 60°C for 20 sec). Control, FASTKD4 KO, and FASTKD4-FLAG rescue RNA utilized for RNA-seq was also used for RT-qPCR.

### Protein stability experiment

To assay the stability of OXPHOS proteins, 1.5x10^5^ NT and FASTKD4 KO HEK293T cells were seeded and the following day were treated with 50 µg/mL chloramphenicol (CAP, Sigma C0378) for 0, 2, or 4 days. On the fifth day after seeding, cells were lysed and OXPHOS protein levels were quantified by immunoblotting as described previously. To determine the change in protein levels over time, the level of protein at 2 or 4 days of CAP treatment was divided by the level of protein at 0 days of CAP treatment.

### Mitoribosome profiling

2.5x10^7^ cells HEK293T NT and FASTKD4 KO were grown under the standard conditions described previously. Media was removed on ice, and cells were rinsed twice with ice-cold PBS. Cells were then scraped into 600 µL of mitoribosome lysis buffer (0.25% lauryl maltoside, 50 mM NH_4_Cl, 20 mM MgCl_2_, 0.5 mM DTT, 10 mM Tris, pH 7.5, and 1× EDTA-free protease inhibitor cocktail (Roche)) and homogenized with six strokes using the DWK 1mL tissue grinder.

Lysates were RNA digested using 8 units/µL of RNaseI (NEB) for 30min at room temperature to generate mitoribosome footprints. The digestion was inhibited by the addition of 80 units SUPERaseIn (Thermo Fisher Scientific), and the lysate was cleared by a 5-minute spin at 10, 000 rpm. To isolate monosomes, 450 µL of clarified lysate was layered onto a 10-50% linear sucrose gradient and centrifuged at 40, 000 rpm for 3 hr at 4°C using an SW41Ti rotor. Gradients were fractioned into 800 µL fractions using the BioComp instrument, and the mitoribosomes were followed using western blots against mitoribosome subunits MRPL12 and MRPL18B. RNA was extracted from the mitomonosome fraction using 1:1 acid phenol/chloroform extraction and resolved on a 15% polyacrylamide TBE-urea gel to collect 28-40 nt RNA fragments. Library preparation was performed as previously described, with modifications specific to mitoribosome profiling protocols (26). The samples were then sequenced on Illumina-based Next-Seq 500 (single-end reads, 75 cycles).

Raw data were processed through a custom bioinformatic pipeline (https://github.com/churchmanlab/human-mitoribosome-profiling). In brief, adapter sequences were trimmed, and filtered reads were aligned to the reference genome using STAR. Mitoribosome footprint periodicity and A-site offsets were established to quantify synthesis rates as transcripts per million (TPM) using featureCounts (Subread). The optimal A-site offset for each mitoribosome profiling library was established by identifying the start codon position for initiating ribosomes on genes with 5’ UTRs (*CO3* and *ND4*) and refined by determining the predominant subcodon position of the 3′ ends for every read length across all coding sequences. We generally applied the following offsets from the 3′ terminus: 32 nt: [−13], 33 nt: [−14], 34 nt: [ −15], 35 nt: [−16], 36 nt: [−17], 37 nt: [ −18], 38 nt: [−19], 39 nt: [−20], 40 nt: [ −20]. Nucleotides that were soft-clipped were excluded from our analysis, except for 3′ adenines that likely represented terminal polyadenylation.

For v-plot analyses, ribosome footprints were assigned to processed or unprocessed classes based on their 5′ endpoint position relative to the annotated mature transcript boundary and, where appropriate, their inferred A-site position. Footprint length was calculated from the mapped 5′ and 3′ endpoints, and analyses were restricted to ribosome-footprint-sized reads, 32–40 nt, unless otherwise indicated. A-site positions were estimated from the 5′ endpoint using a length-dependent offset. Processed start-like footprints were defined as reads whose 5′ ends mapped near the mature processed transcript boundary, generally within −2 to +2 nt of the annotated start, and whose inferred A-sites fell in the expected early coding-region window downstream of the start site. For the *ATP8/6-CO3* and *ND5-CYB* junctions, this processed A-site window was defined as +15 to +23 nt relative to the processed start. These reads were interpreted as ribosomes initiating or early elongating on the mature processed downstream mRNA. Populations of reads longer than > 40 nt, that displayed prominent accumulation near the start site but did not exhibit the coordinated 5′-and 3′-end characteristics of translating ribosomes were excluded from start-site classification.

Translational efficiency for mitochondrial-encoded transcripts was estimated by dividing ribosome-protected footprint counts from ribosome profiling in transcripts per million (TPM) by RNA abundance. Matched ribosome profiling and RNA-seq libraries were not available, so translational efficiency was approximated by dividing the ribosome profiling-derived TPM for a given gene by the DESeq2 median-of-ratios-normalized gene counts generated from the RNA-sequencing experiment described above.

### DMS-MaPseq

#### In organello DMS labeling

Mitochondrial DMS-MaPseq was performed as described in Moran et al. (27). NT and FASTKD4 KO HEK293T cells were grown to 80-90% confluency in 500 cm^2^ square TC-treated culture dishes (Corning) in DMEM supplemented with 10% FBS. Mitochondria were isolated as described previously and resuspended in 1 mL STE buffer (0.32 M sucrose, 1 mM EDTA, 10 mM Tris-HCl pH 7.5). For each replicate, 4.5 mg of freshly prepared mitochondria were pelleted at 10, 000*g* for 10 min, and the pellet was resuspended in 225 µL Na-cacodylate buffer (300 mM Na-cacodylate, 6 mM MgCl_2_, 0.32 M sucrose) that was preheated to 37°C. 225 µL of preheated Na-cacodylate buffer plus 4.5 µL of DMS was then added to the mitochondria solution to achieve a 1% DMS concentration. DMS-mitochondria solution was incubated at 37°C for 5 min, with gentle mixing by inverting every minute. The reaction was quenched by the addition of 270 µL β-mercaptoethanol. Samples were centrifuged at 10, 000*g* for 10 min to pellet the mitochondria. The mitochondrial pellet was washed twice in 1 mL of glycerol buffer (10% glycerol, 0.15 mM MgCl_2_, 10 mM Tris-HCl pH 6.8) before resuspension in Trizol. RNA was extracted from the mitochondria following manufacturer’s instructions and precipitated using 100% isopropanol and 3 µL glycogen.

#### rRNA depletion

10 µg of mitochondrial RNA was DNase treated with the TURBO DNase kit (ThermoFisher Scientific) and purified using the RNA Clean and Concentrator -5 kit (Zymo). A custom rRNA depletion protocol, established by Moran et al., was used to subtract cytosolic and mitochondrial rRNAs from the samples (27). In brief, a 10x mix of oligomers against mitochondrial and cytosolic rRNAs was prepared at a 2:1 mitochondrial:cytosolic ratio, to give a final concentration of 500 nM per mitochondrial oligo and 250 nM per cytosolic oligo. 1 µg of DNase-treated RNA was used as the input for each rRNA depletion reaction, with multiple depletion reactions being performed for each sample to achieve sufficient input for subsequent library preparation. 1 µL of the 10x oligo mix and 2 µL of 5x hybridization buffer (1 M NaCl, 0.5 M Tris-HCl pH 7.5) were added to each depletion reaction and the final volume was adjusted to 10 µL with RNase-free water. Samples were denatured at 95°C for 2 min before the temperature was reduced 0.1°C/sec down to 22°C. 2 µL RNase H buffer, 2 µL thermostable RNase H, and 6 µL water were added to the samples which were then incubated at 65°C for 5 min. The rRNA depletion oligos were degraded by DNase treatment using the TURBO DNase kit and RNA was cleaned with RNA Clean and Concentrator -5 kit following manufacturer’s protocol to select RNAs >200 nt in size.

#### Library preparation

The strand-aware xGen Broad-range RNA library preparation kit (IDT) was utilized to generate sequencing libraries following manufacturer’s instructions with the modifications described in Moran et al. (27). In brief, the RNA fragmentation step was performed by combining 50 ng of rRNA depleted RNA with 1 µL reagent F1, 4 µL reagent F3, and 2 µL F4, followed by incubation at 95°C for 2 min. Samples were then immediately placed on ice and combined with 1 µL of Induro reverse transcriptase (New England Biolabs), 1 µL reagent R1, and 1 µL 0.1 M DTT. The RNA:reverse transcriptase solution was incubated at room temperature for 30 min before the addition of 2 µL reagent F2, and then the solution was placed in a thermocycler for 10 min at 20°C, 10 min at 42°C, and 60 min at 55°C. A pause was programmed into the thermocycler protocol at 55°C, during which 1 µL of 4 M NaOH was added before the program continued for 3 min at 95°C. The sample was then removed from the thermocycler and placed on ice and 1 µL of 4 M HCl and 27 µL of low EDTA TE were added. Following the indexing PCR, using 12 PCR cycles, indexed libraries were resolved on an 8% TBE gel for 55 min at 180 V to size select for products ∼300 nt. These products were then sequenced with NovaSeqX system performing a paired-end run for 150 cycles.

### DMS-MaPseq data analysis and secondary structure prediction

DMS mutational profiling data were processed using the SEISMIC workflow (https://github.com/rouskinlab/seismic-rna) to generate nucleotide-resolution mutation-rate profiles. Briefly, sequencing reads were aligned to the mitochondrial reference sequence, and nucleotide substitutions were identified relative to the reference. SEISMIC was then used to calculate the mutation rate at each nucleotide position from the fraction of informative reads containing a mutation at that position. Nucleotide positions with mutation rates greater than 15% were excluded from the folding analysis. The filtered nucleotide-resolution mutation rates were used as experimental constraints for RNA secondary-structure prediction with RNAstructure (28). Folding was performed independently for each 250-nt region using the corresponding mitochondrial reference sequence and DMS-derived mutation profile. The minimum-free-energy structure predicted for each region was retained for downstream visualization and interpretation. Predicted RNA structures were visualized using VARNA (29). Nucleotides were colored according to their SEISMIC-derived mutation rates using discrete reactivity categories.

### Statistics and reproducibility

All statistical analyses were performed using GraphPad Prism or R software. No statistical methods were used to predetermine sample size. No data were excluded from analysis. Experiments were not randomized and data collection and analysis were not performed blind to the experimental conditions. Data met the assumptions of the statistical tests indicated.

## Results

### FASTKD4 loss selectively alters mitochondrial RNA and protein expression

To examine how non-canonical processing affects mitochondrial gene expression, we generated three independent CRISPR/Cas9 FASTKD4-knockout (KO) clones in HEK293T cells (Fig. S1A). We observed that FASTKD4-deficient cells exhibited reduced ATP production under basal glucose conditions and when glycolysis was inhibited by 2-deoxy-D-glucose (2-DG) compared to the non-targeting (NT) control, consistent with an impaired capacity for oxidative phosphorylation (OXPHOS; Fig. S1B). However, these cells did not display elevated apoptosis under basal or glycolysis-inhibited conditions (Fig. S1C).

Immunoblotting revealed a significant decrease in CYB and COX1 protein levels in FASTKD4-deficient cells, whereas ND1 and ATP6 were unchanged (Fig. 1B, S1D). RNA-sequencing demonstrated that, while the steady-state abundance of *ATP6*, *CO3*, and *ND3* was reduced, there was no change in *CYB* transcript levels that could explain the depletion of CYB protein (Fig. 1C). CYB protein stability was also unaffected by loss of FASTKD4 (Fig. S1E). The expression phenotypes observed in FASTKD4-depleted cells were rescued by ectopic expression of FLAG-tagged FASTKD4 and were reproduced across orthogonal assays and cell lines (Fig. 1D, S1F-H). These results reveal the transcript-specific effects of FASTKD4 loss and raise the question of how defective processing alters distinct stages of mitochondrial gene expression.

### FASTKD4 promotes mtRNA processing at non-canonical junctions

We hypothesized that the differential effects on mitochondrial gene products in FASTKD4-deficient cells resulted from the retention of unprocessed precursor transcripts at non-canonical junctions. To directly assess RNA processing states in FASTKD4-depleted cells, we used Oxford Nanopore long-read RNA sequencing to quantify individual mitochondrial transcripts retaining unprocessed 5′ or 3′ junctions. For genes flanked by tRNAs, such as *ND1, ND2, CO2, ND3*, the fractions of unprocessed 5′ and 3′ junctions were low and changed only modestly upon loss of FASTKD4 (Fig. 2A-B). In contrast, non-canonical junctions displayed a pronounced accumulation of unprocessed ends in FASTKD4-deficient cells. Within the subset of non-canonical junctions, loss of FASTKD4 impaired the processing of the *ND5-CYB* junction with a drastic increase in 5′ unprocessed *CYB* reads accompanied by a corresponding increase in 3′ unprocessed *ND5* reads (Fig. 2A-B). These data implicate FASTKD4 as a crucial factor involved in the processing of the *ND5-CYB* precursor. We also detected elevated unprocessed ends at the *ATP8/6-CO3* non-canonical junction. In contrast, we did not observe large changes in the fraction of processed ends for the non-canonical 5′ *CO1* junction upon FASTKD4 loss (Fig. 2A). Collectively, these single-molecule long-read mt-mRNA sequencing data demonstrate that loss of FASTKD4 impairs the processing of mt-mRNAs specifically at non-canonical junctions, while processing of canonical, tRNA-flanked junctions remains largely intact.

**Figure 2:**
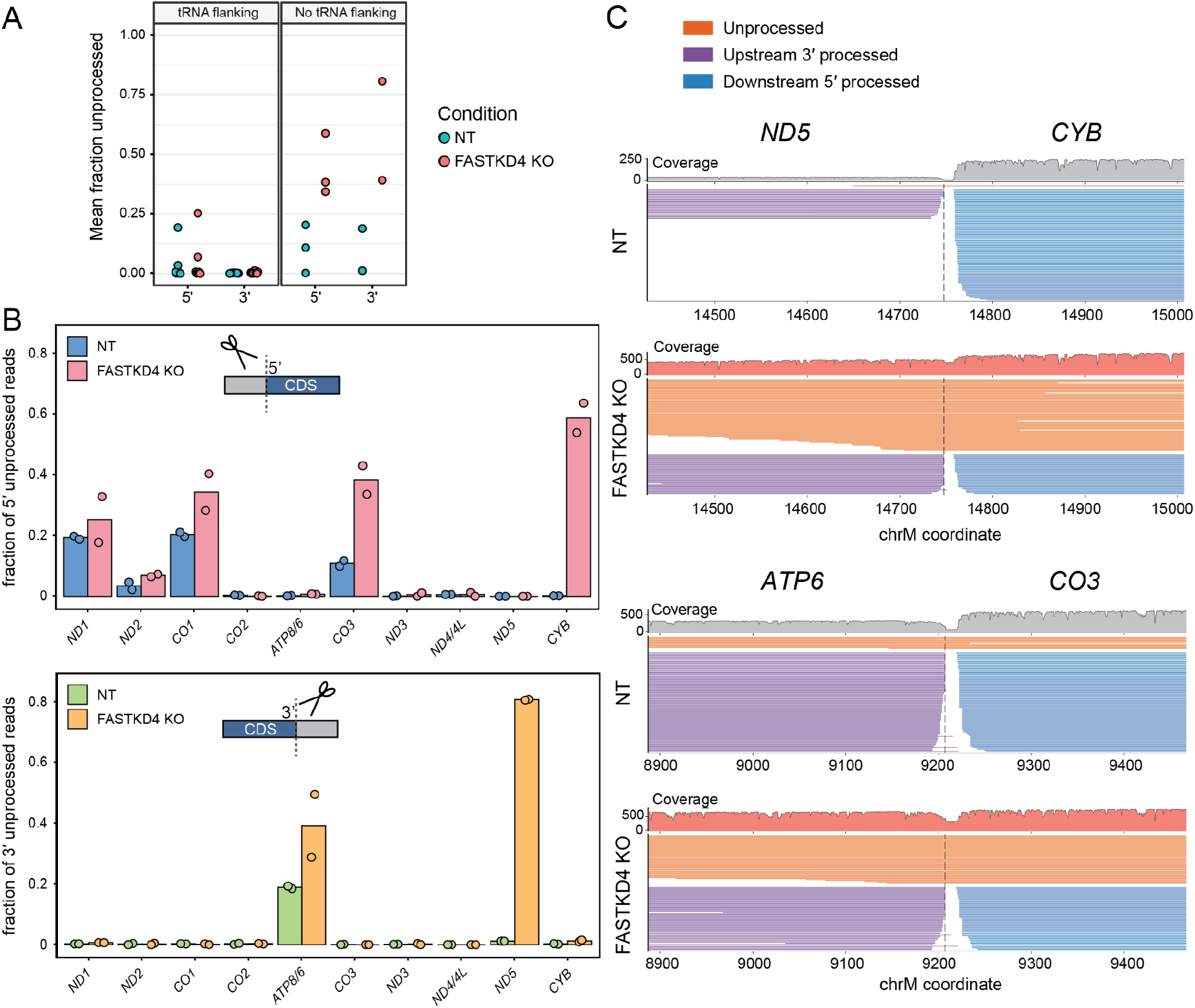
FASTKD4 depletion impairs mt-mRNA processing at non-canonical junctions. **A)** Mean fraction 5′ and 3′ unprocessed transcripts measured by direct RNA sequencing for genes flanked by a tRNA or lacking a flanking tRNA in NT and FASTKD4 KO cells. **B)** Fraction of 5′ (top) and 3′ (bottom) unprocessed reads from Nanopore direct RNA sequencing for heavy strand mt-mRNAs from two replicate measurements of total RNA from NT and FASTKD4 KO cells. **C)** Gene track depicting a subsampling of direct RNA sequencing reads aligned to the *ND5-CYB* (top) and *ATP8/6-CO3* (bottom) junctions in NT and FASTKD4 KO cells. Scale indicates read coverage represented by the height of the track.

### CYB protein depletion is not explained by altered transcript turnover

Steady-state RNA abundance does not capture transcript turnover, which determines how long an individual transcript is available to be engaged by ribosomes and support protein synthesis. We therefore asked whether altered *CYB* stability could contribute to reduced CYB protein levels in FASTKD4-deficient cells. As mt-mRNAs are transcribed on a single polycistronic precursor, differences in steady-state abundance are largely determined by transcript-specific degradation rates (19, 30). To measure mtRNA turnover, we labeled cells with 4-thiouridine (4sU) for 90 minutes, biotinylated and depleted newly synthesized 4sU-containing RNA, and quantified the remaining “old”, unlabeled RNA using MitoStrings (Fig. S2A; (19, 31, 32)). Steady-state transcript abundances measured by MitoStrings were largely consistent with our RNA-sequencing data (Fig. S2B). FASTKD4 depletion increased the turnover of *ATP8/6* and *CO3* and decreased the turnover of *ND1* and *ND5* (Fig. S2C). By contrast, *CYB* turnover was unchanged, excluding reduced transcript lifetime as the cause of CYB protein depletion.

In addition, loss of FASTKD4 moderately dysregulated poly(A) tail length for a subset of transcripts, including *ND1*, *CO2*, and *ND3*, but these changes did not consistently correlate with transcript turnover in our measurements (Fig. S2D). Shortening of the *ND3* poly(A) tail in FASTKD4-deficient cells did coincide with a dramatic reduction in *ND3* steady-state abundance, as previously described (14). We noted that, unlike most mt-mRNAs, *ND3* steady-state abundance depended on FASTKD4 rather than LRPPRC, a broad regulator of mt-mRNA stability (Fig. S2E-F; (14, 19, 31–33)). These findings point to additional transcript-specific roles for FASTKD4 in mtRNA metabolism beyond non-canonical junction processing.

### FASTKD4 loss selectively reduces *CYB* translational efficiency

Having established that neither the steady-state abundance nor turnover of *CYB* was altered in FASTKD4-deficient cells, we asked whether reduced translation could account for the depletion of CYB protein. Sucrose gradient fractionation revealed no defects in mitoribosome assembly in FASTKD4-deficient cells, as compared to control cells (Fig. S3A). We therefore used mitoribosome profiling to directly assess translation across the mitochondrial translatome (34, 35). Quantification of mitoribosome-protected footprints (RPFs) mapping to each mt-mRNA as transcripts per million (TPM) revealed reduced RPFs mapping to *CO3*, *ND3*, and *CYB* in FASTKD4-depleted cells compared to the control, while RPFs for several other transcripts, such as *CO2* and *ND4L* were increased, consistent with previous observations of altered protein synthesis rates in FASTKD4-depleted cells (Fig. S3B; (13)).

The total transcript abundance of *CO3* and *ND3* was reduced in cells lacking FASTKD4, thus the decrease in their RPFs could reflect fewer available transcripts rather than a translational defect. To distinguish between these possibilities, we estimated translational efficiency for each transcript by normalizing RPF counts to steady-state RNA abundance (Fig. 3A). Indeed, the translational efficiencies of *CO3* and *ND3* were unchanged between FASTKD4-depleted cells and the control (Fig. 3B). In contrast, the translational efficiency of *CYB* was substantially reduced in FASTKD4-depleted cells. This selectivity was unexpected: impaired processing of both the *ATP8/6-CO3* and *ND5-CYB* junctions resulted in strongly reduced translational efficiency of *CYB*, but not *CO3*. The observation that two similarly misprocessed junctions yield distinct translational outcomes suggests that features intrinsic to the junctions themselves may determine whether unprocessed transcripts remain competent for translation.

**Figure 3:**
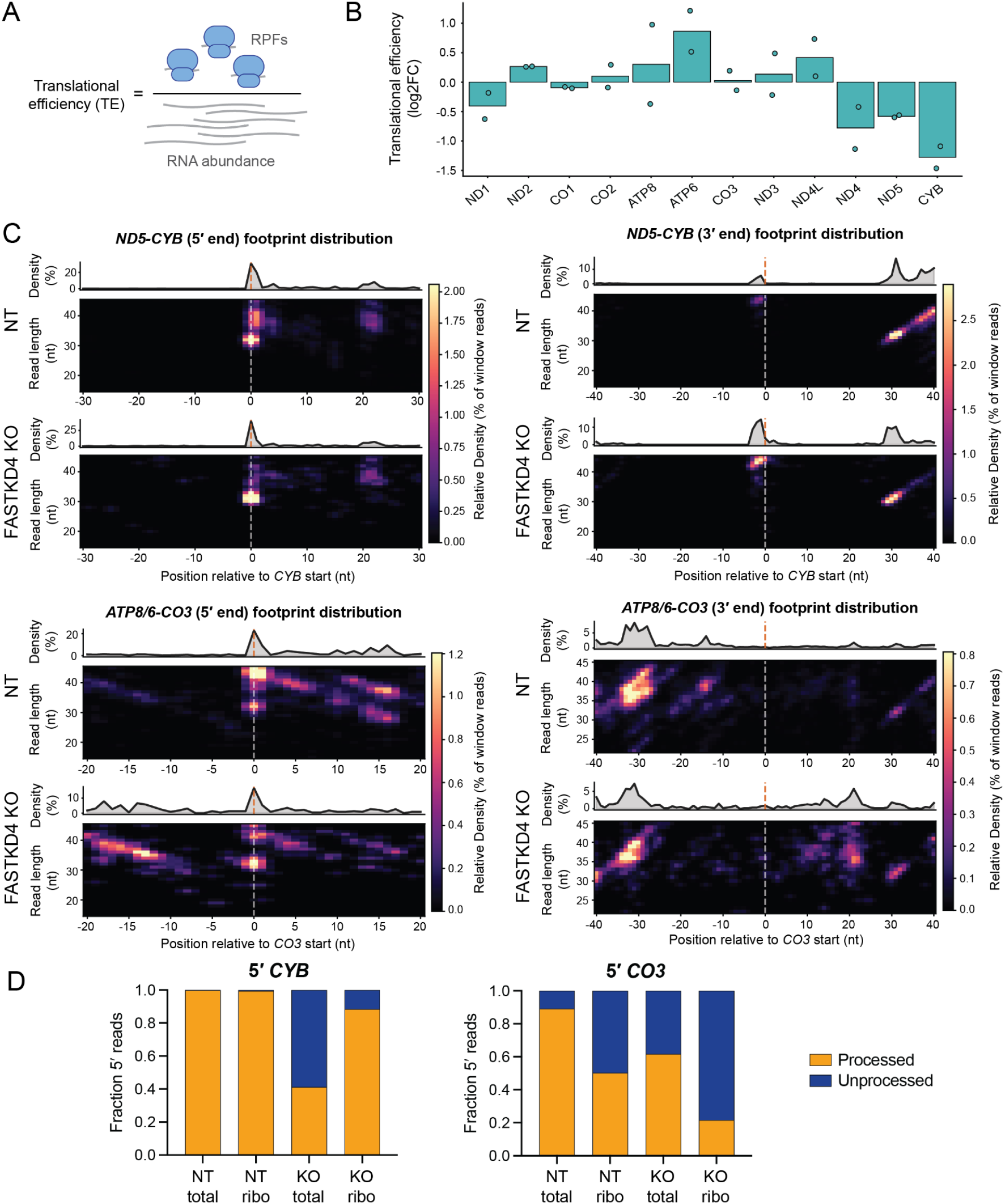
Reduced processing of *ND5-CYB* junction in FASTKD4 KO cells impairs translational efficiency of *CYB*. **A)** Translation efficiency (TE) was calculated by dividing TPM values determined by mitoribosome profiling by RNA abundance values determined by RNA-sequencing for each transcript. **B)** Log2FC (FASTKD4 KO/NT) in TE for each heavy-strand transcript (right). TPM, transcripts per million. **C)** Heatmap depicting the 5′ (left) and 3′ (right) ends of mitoribosome profiling reads aligned to the *ND5-CYB* junction (top) and *ATP8/6-CO3* (bottom) in NT and FASTKD4 KO cells. Dashed line represents the annotated 5′ end of the mature *CO3* or *CYB* transcript. **D)** Fraction 5′ processed transcripts from total RNA (total) and ribosome-associated RNA (ribo) in NT and FASTKD4 KO cells for *CYB* (left) and *CO3* (right).

### Mitoribosomes preferentially engage processed *CYB* transcripts

The reduced translational efficiency of *CYB* in FASTKD4-deficient cells, despite unchanged transcript abundance, suggested that the unprocessed *ND5-CYB* precursor may be a poor substrate for the mitoribosome. To test this, we analyzed our mitoribosome profiling data to determine whether RPFs mapped to processed or unprocessed transcripts. We quantified the fraction of RPFs that spanned beyond the annotated 5′ or 3′ gene ends as a measure of ribosomal density on unprocessed molecules. For most genes, in both control and FASTKD4-depleted cells, over 90% of footprints mapped to processed transcripts (Fig. S4A). The exception to this trend was *ATP8/6-CO3*. About 60% of RPFs mapped to processed *CO3* transcripts in control cells and ∼20% in FASTKD4-deficient cells, suggesting that the mitoribosome loads onto unprocessed *ATP8/6-CO3* transcripts, consistent with prior studies (19, 26, 36).

Analysis of RPF end positions across footprint lengths revealed distinct patterns of mitoribosome occupancy at the *ND5-CYB* and *ATP8/6-CO3* junctions. (Fig. 3C). At the *ND5-CYB* junction, RPF positioning was similar in control and FASTKD4-deficient cells despite the pronounced accumulation of unprocessed precursor in the latter. Most RPF 5′ ends aligned with the annotated *CYB* 5′ boundary, with a small fraction extending upstream, while the corresponding 3′ ends mapped into the *CYB* coding region in a footprint-length-dependent manner. This geometry is consistent with ribosomes positioned on leaderless, processed *CYB* transcripts and indicates little engagement of the unprocessed precursor (26, 37). In contrast, at the *ATP8/6-CO3* junction, RPF 5′ ends mapped both to the annotated *CO3* start and upstream within the *ATP6* coding region. The corresponding 3′-end distributions and classification of junction-associated footprints supported mitoribosome occupancy on both processed and unprocessed *CO3* transcripts in control and FASTKD4-deficient cells (Fig. S4B, S5A-B). Thus, mitoribosomes preferentially engage processed *CYB* transcripts but can occupy both processed and unprocessed *CO3* transcripts.

Comparing the fraction of unprocessed transcripts in the total RNA pool and in the mitoribosome-protected footprints revealed junction-specific selectivity in ribosome engagement. Although 5′ unprocessed *CYB* transcripts accumulated in FASTKD4-depleted cells, only a small fraction of *CYB*-associated mitoribosome footprints mapped to the unprocessed precursor (Fig. 3D). In contrast, a significant proportion of 5′ unprocessed *CO3* transcripts were engaged by the mitoribosome upon loss of FASTKD4. Together, these data suggest that mitoribosomes largely exclude unprocessed *ND5-CYB* precursors yet engage unprocessed *ATP8/6-CO3* transcripts, pointing to a specific feature of the *ND5-CYB* junction that impedes translation initiation.

### RNA secondary structure within the *ND5-CYB* junction may impede *CYB* translation

After processing from the polycistronic precursor, the majority of mature mt-mRNAs are leaderless, lacking a 5′ UTR preceding the start codon (38, 39). The 5′ regions of these leaderless transcripts have been shown to be unstructured both *in vitro* and *in vivo*, a feature proposed to facilitate mitoribosome loading (27, 40). When processing of the *ATP8/6-CO3* and *ND5-CYB* junctions is impaired, both *CO3* and *CYB* retain an aberrant 5′ extension. Given that *CYB* translation is decreased in FASTKD4-depleted cells, we hypothesized that this extension introduces RNA secondary structure at one junction but not the other, explaining the selective translational defect on unprocessed *CYB*.

To test this, we used mitochondrial DMS mutational profiling with sequencing (mitoDMS-MaPseq) to probe RNA structure across the mitochondrial transcriptome in isolated mitochondria (27). Dimethyl sulfate (DMS) methylates the base-pairing faces of accessible adenines and cytosines, so measures of DMS reactivity are indicative of base accessibility. DMS reactivities were well-correlated between replicates and similar between FASTKD4-deficient and control cells, indicating that loss of FASTKD4 does not broadly alter base accessibility across the mitochondrial transcriptome (Fig. S6A-B). We used our experimentally determined DMS reactivities as input for RNA folding algorithms to predict secondary structures at the two non-canonical junctions (28, 41).

Concordance with prior transcriptome-wide mitochondrial RNA structure inference benchmarked our approach and supported its use for identifying junction-specific structural features (Fig. S6C, S7; ((27)). The majority of mt-mRNAs exhibited moderate to high DMS reactivity near their start codons, within their first 50 nucleotides, and were largely predicted to be single stranded, consistent with previous reports (Fig. S7; (27, 40). Predicted secondary structure across the processed *CYB* transcript was largely preserved in FASTKD4-deficient cells, with linear arc plots revealing only localized differences in base-pairing between knockout and control cells (Fig. 4A). Notably, the 5′ processed *CYB* transcript was predicted to remain unstructured around the start codon, consistent with its preferential engagement by the mitoribosome.

**Figure 4:**
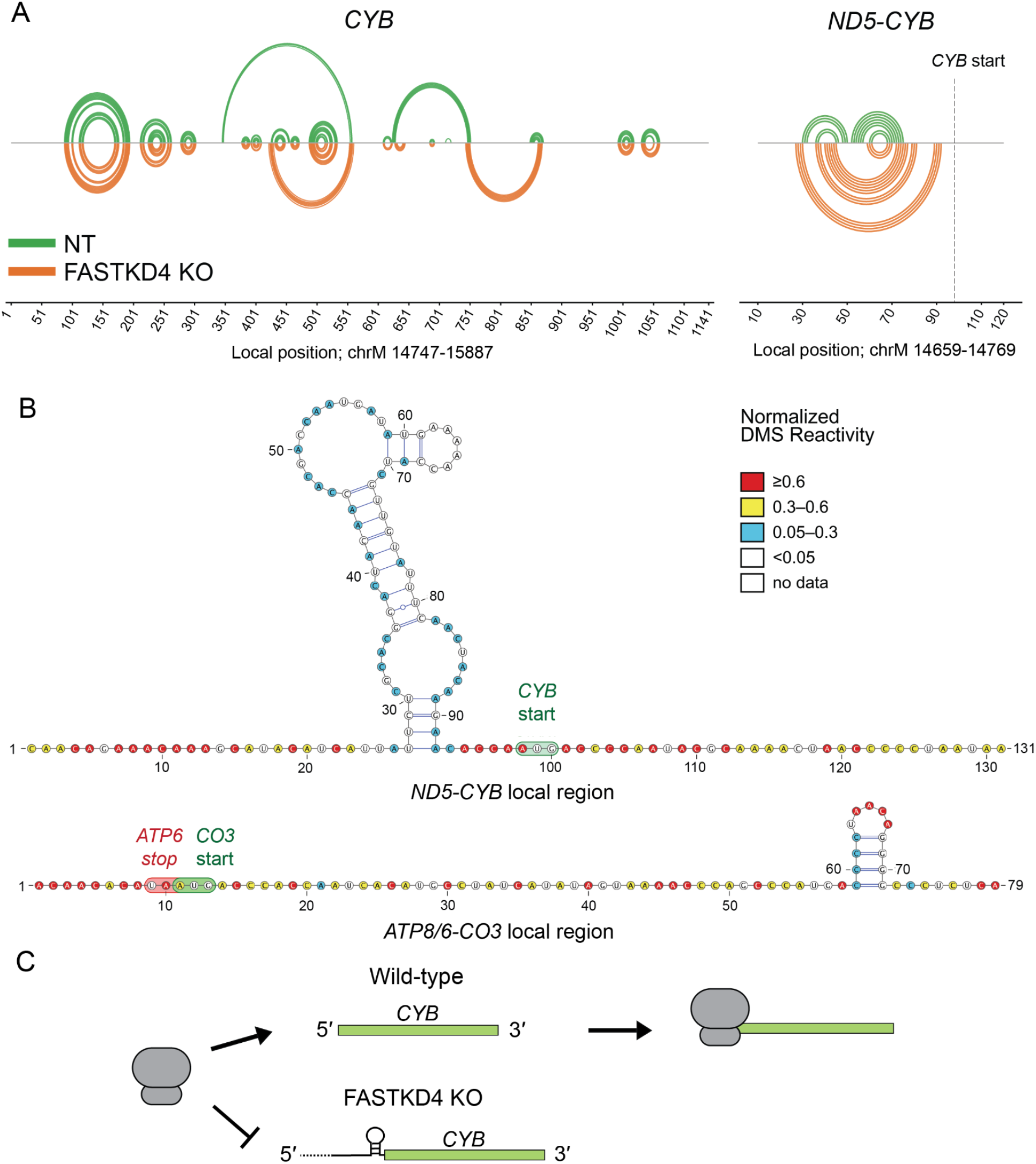
RNA secondary structure within the *ND5-CYB* junction may impede *CYB* translation. **A)** Linear arc plot depicting changes in predicted RNA secondary structure between NT and FASTKD4 KO for *CYB* (left) and the *ND5-CYB* junction (right). **B)** Predicted secondary structure of unprocessed *ND5-CYB* (top) and *ATP8/6-CO3* (bottom) junctions in FASTKD4 KO. Nucleotides colored by normalized DMS reactivity according to the scale shown. Start and stop codons are highlighted. DMS, dimethyl sulfate. **C)** Schematic of structure-dependent mitoribosome engagement with *CYB* in wild-type and FASTKD4-depleted cells.

We then compared the predicted secondary structures within the *ATP8/6-CO3* and *ND5-CYB* junctions in FASTKD4-deficient cells, where the unprocessed precursor accumulates. At the *ND5-CYB* junction, several nucleotides upstream of the *CYB* start codon displayed low DMS reactivity in FASTKD4-deficient cells. DMS reactivity-constrained folding predicted a possible secondary structure located just five nucleotides upstream of the *CYB* start codon (Fig. 4A-B). By contrast, most nucleotides at the *ATP8/6-CO3* junction were highly DMS-reactive, and the region surrounding the *CO3* start codon was predicted to remain unstructured (Fig. 4B). These contrasting structural features mirror the distinct translational outcomes at the two junctions: mitoribosomes engage unprocessed *CO3* transcripts and *CO3* translational efficiency is maintained, whereas *CYB* translational efficiency is selectively impaired.

Together, these findings support a structural explanation for the junction-specific translational effects of FASTKD4 loss. Retention of the *ND5-CYB* junction permits base-pairing immediately upstream of the *CYB* start codon, which may impede translation initiation. By contrast, the *CO3* start codon remains accessible within the unprocessed *ATP8/6-CO3* precursor. Thus, non-canonical processing is not universally required for translation, but may license *CYB* translation by removing structural elements near its start codon with the precursor RNA (Fig. 4C).

## Discussion

Non-canonical processing is not uniformly required for mitochondrial translation. Although FASTKD4 loss impaired processing at both the *ND5-CYB* and *ATP8/6-CO3* junctions, only *CYB* translational efficiency declined, resulting in reduced CYB protein abundance. Mitoribosomes engaged 5′ unprocessed *CO3*, but largely excluded 5′ unprocessed *CYB* transcripts. DMS reactivity-constrained folding suggests a basis for this selectivity: a predicted structural element lies adjacent to the *CYB* start codon in the unprocessed *ND5-CYB* precursor, while the corresponding region surrounding the *CO3* start codon is predicted to remain linear and accessible. This contrast shows that retention of a 5′ extension is not intrinsically incompatible with human mitochondrial translation. Instead, local sequence and structural features determine whether a 5′ unprocessed transcript remains translation competent. Thus, non-canonical processing not only defines mature RNA ends. At select junctions, it converts a poorly translated precursor into a translation-competent mRNA, coupling RNA maturation to protein output.

The mechanism by which FASTKD4 promotes non-canonical processing remains unresolved. Structural modeling suggests that FASTKD4 resembles the bacterial Vsr endonuclease and could cleave RNA directly; however, endonuclease activity has not been demonstrated (11, 14). Supporting the possibility that FASTK proteins can act directly in junction processing, FASTKD5 was recently shown to cleave RNA substrates (15). While loss of either FASTKD4 and FASTKD4 impairs non-canonical junction processing, *in vitro* and *in vivo* comparisons indicate that FASTKD4 and FASTKD5 have distinct junction preferences: FASTKD4 more strongly promotes *ND5-CYB* processing, whereas FASTKD5 more strongly promotes processing at the 5′ end of *CO1* (11, 13, 15). Consistent with this division of activity, we found that FASTKD4 loss impaired *ND5-CYB* processing, but had minimal effect on 5′ *CO1* processing. Although COX1 protein levels were moderately reduced, this may be an indirect consequence of decreased *CO3* expression and impaired complex IV assembly rather than defective *CO1* RNA processing.

We demonstrated that loss of FASTKD4 selectively reduced the translational efficiency of *CYB*. The reduced RPFs mapping to *CO3* and *ND3* in FASTKD4-depleted cells were accounted for by their decreased transcript abundance, and the translational efficiencies of these transcripts were unchanged. In contrast, the translational efficiency of *CYB* was substantially reduced despite unchanged transcript levels. This selectivity raised the question of what distinguishes the *ND5-CYB* junction from the *ATP8/6-CO3* junction at the molecular level.

*In vitro,* mitoribosomes preferentially load on 5′ leaderless mRNAs, and impaired processing of the 5′ *CO1* non-canonical junction has been shown to reduce ribosome binding and COX1 protein synthesis *in vivo* (13, 15, 19, 42). These observations might suggest that any 5′ extension would be deleterious for mitoribosome loading. However, the mitoribosome readily engages unprocessed *CO3* transcripts bearing a 5′ extension, and is known to translate internal start codons on the bicistronic *ATP8/6* and *ND4L/4* transcripts (19, 26, 36). Thus, the presence of upstream sequence alone is not sufficient to block translation, and the selective exclusion of unprocessed *CYB* from the mitoribosome points to a feature specific to the *ND5-CYB* junction.

RNA secondary structure in the 5′ UTR is a well-characterized determinant of translation efficiency in both eukaryotic cytosolic and bacterial systems (43–45). In the mitochondria, the 5′ regions of mature leaderless mt-mRNAs are unstructured both *in vitro* and *in vivo*, which could facilitate mitoribosome loading (27, 40). Our DMS-MaPseq data extend this principle to the context of failed non-canonical processing: the unprocessed *ND5-CYB* precursor contains a predicted secondary structure adjacent to the *CYB* start codon, while the *ATP8/6-CO3* junction remains unstructured. Thus, non-canonical processing effectively restores the leaderless, unstructured 5′ end that the mitoribosome favors for efficient translation. When processing fails at the *ND5-CYB* junction, the retained upstream sequence introduces a structural barrier adjacent to the start codon. In contrast, at the *ATP8/6-CO3* junction, the equivalent upstream sequence is unstructured and translation proceeds. More broadly, these findings suggest that cis-elements within precursor RNAs can impede translation initiation, positioning non-canonical processing and its associated factors as regulators of mitochondrial translation.

In conclusion, our results reveal that non-canonical mtRNA processing promotes translation at select junctions by eliminating structural barriers within the immature transcript. These data establish a direct link between mtRNA processing and translational regulation and highlight non-canonical processing as a crucial regulatory step in the mt-mRNA lifecycle. An important open question is whether processing at non-canonical junctions is regulated. Mitochondrial stress can alter mtRNA processing and translation, raising the possibility that selective processing of the *ND5-CYB* and *ATP8/6-CO3* junctions could tune complex III and complex IV biogenesis (46). Conservation of the predicted structural element at the *ND5-CYB* junction would further support an evolved role for processing-dependent control of *CYB* translation. Resolving whether FASTKD4 itself possesses the endonucleolytic activity necessary for maturation of non-canonical transcripts will also be essential for understanding this pathway. Furthering our understanding of how cis-elements within precursor transcripts influence mitochondrial gene expression may inform the study of mitochondrial diseases associated with defective RNA processing.

---

**Supplementary Figure 1:**
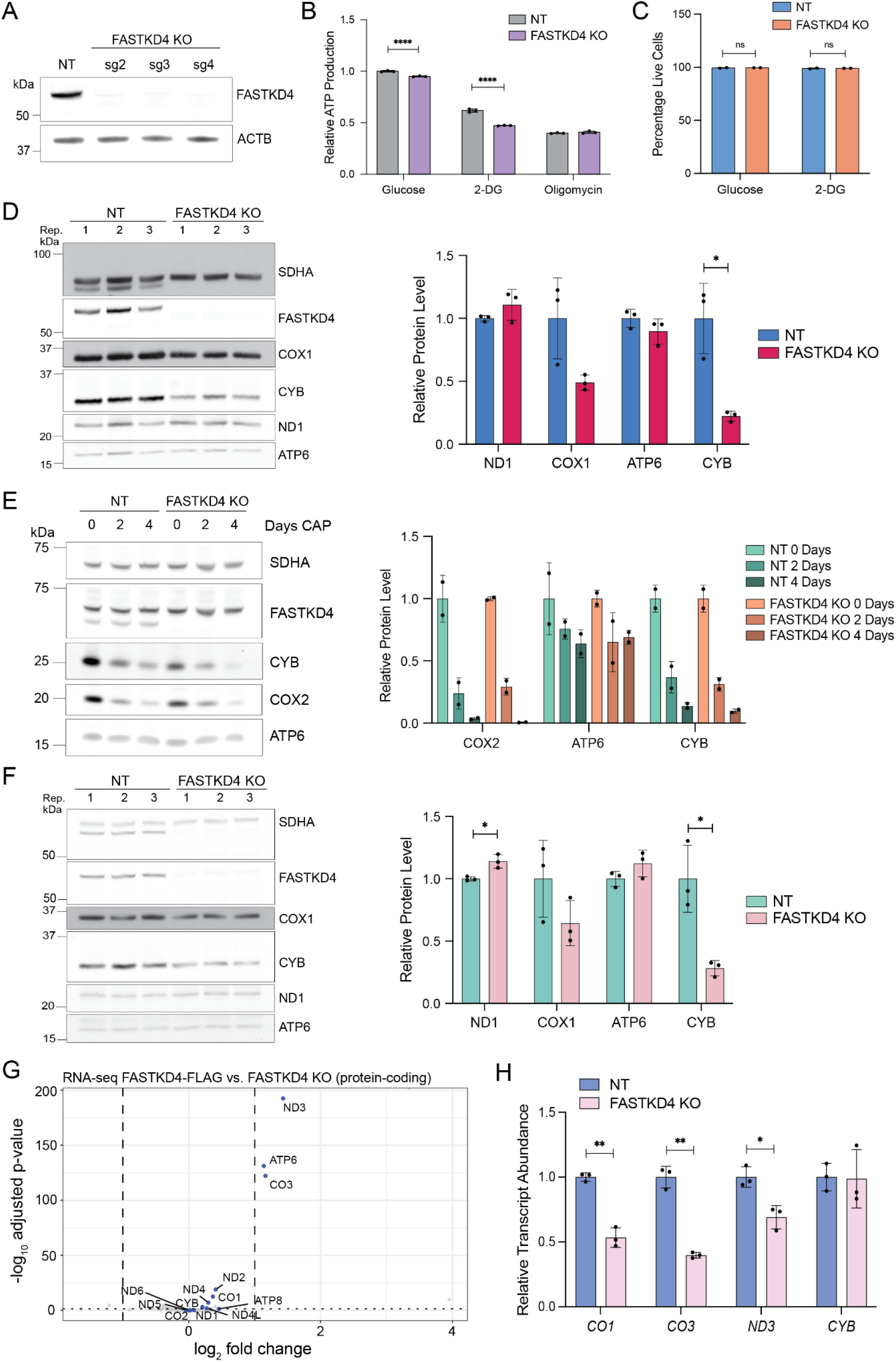
Characterization of FASTKD4 KO in HEK293T and HeLa cells. **A)** Western blot analysis confirming successful FASTKD4 KO in HEK293T cells for three independently generated clonal KO lines. NT, non-targeting control. **B)** CellTiter-Glo assay to quantify relative ATP production by NT and FASTKD4 KO cells when cultured in glucose media compared to media containing 2-DG or oligomycin. Error bars represent standard deviation. 2-DG, deoxyglucose. Two-way ANOVA followed by Sidak’s multiple comparison test, and significant p-values are indicated; ****p-value < 0.0001. **C)** Percentage live cells for NT and FASTKD4 KO cells cultured in either glucose media or media containing 10 mM 2-DG for 4 days, as determined by AnnexinV/PI staining and flow cytometry analysis. Two-way ANOVA followed by Sidak’s multiple comparison test; ns, not statistically significant. **D)** Representative Western blot analysis of OXPHOS subunits (left) and quantification of protein levels (right) from three replicate measurements of whole cell lysate from NT and FASTKD4 KO cells. Error bars represent standard deviation. Unpaired t-test with Welch’s correction, and significant p-values are indicated; *p-value < 0.05. **E)** Representative Western blot of OXPHOS protein stability (left) and quantification of protein levels (right) from two replicate measurements of whole cell lysate from NT and FASTKD4 KO cells after 0, 2, or 4 days of treatment with chloramphenicol (CAP). Error bars show standard deviation. Two-way ANOVA followed by Sidak’s multiple comparison test. Interaction not statistically significant unless otherwise indicated. **F)** Representative Western blot analysis of OXPHOS subunits (left) and quantification of protein levels (right) from three replicate measurements of whole cell lysate from NT and pooled FASTKD4 KO in HeLa. Error bars represent standard deviation. Statistical testing is the same as in **D**. **G)** Volcano plot depicting log2 fold change (FASTKD4-FLAG rescue/FASTKD4 KO; x-axis) and -log10 adjusted p-value (y-axis) from RNA-seq. mt-mRNAs are highlighted. **H)** NT-normalized transcript abundance (relative to ACTB) for a subset of mt-mRNAs as quantified by RT-qPCR for three replicate measurements from NT and pooled FASTKD4 KO HeLa cells. Error bars represent standard deviation. Statistical testing is the same as in **D**; *p-value < 0.05, **p-value < 0.01.

**Supplementary Figure 2:**
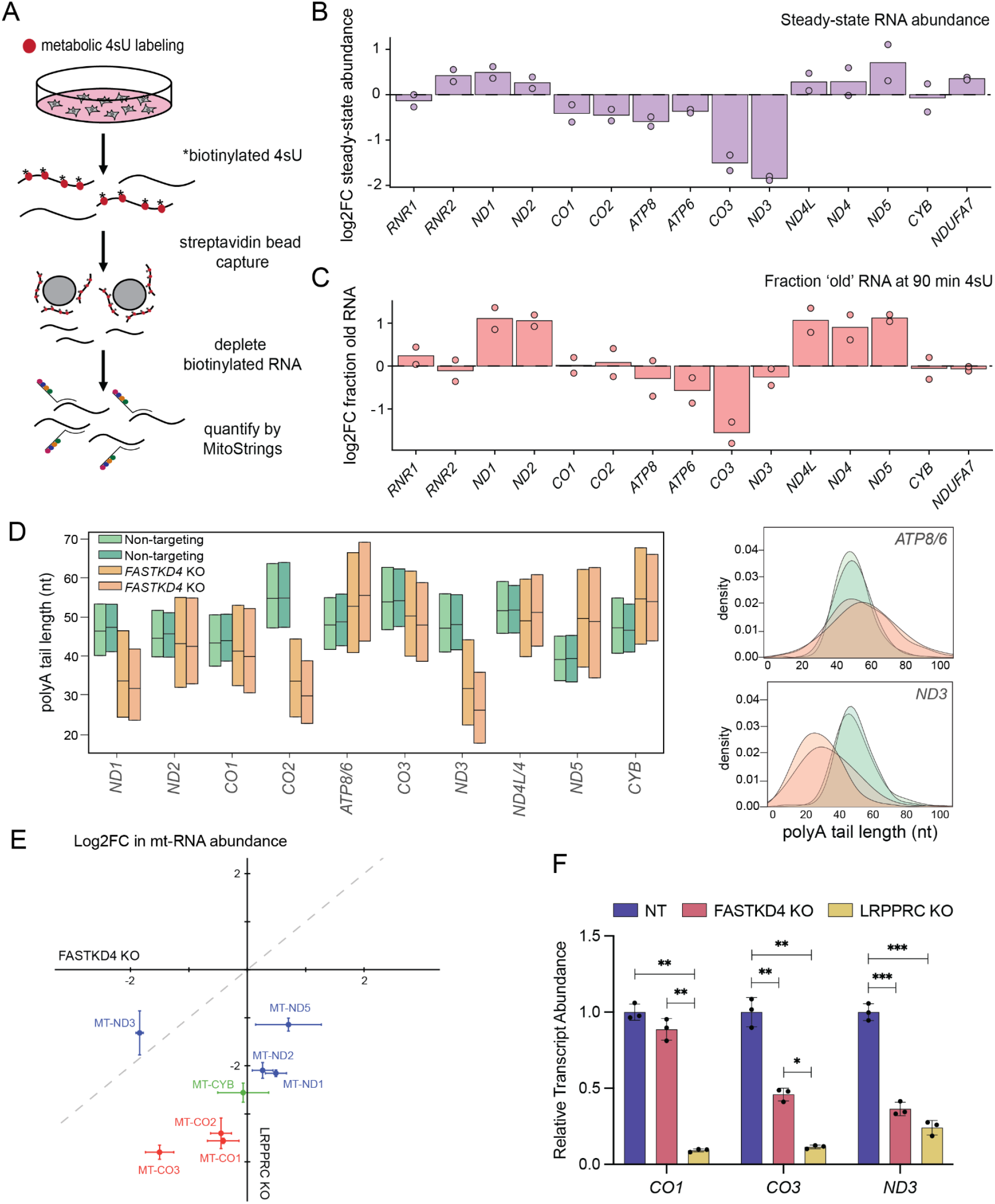
Stability of CYB transcripts is independent of FASTKD4. **A)** Schematic of metabolic labeling of nascent transcripts with 4sU. Cells are treated with 4sU, which is incorporated into nascently transcribed RNA. 4sU is biotinylated and nascent biotinylated RNAs are captured and depleted with streptavidin beads. Unlabeled ‘old’ RNA is quantified by MitoStrings. 4sU, 4-thiouridine. **B)** Log2 fold change in steady-state transcript abundance mtRNAs and nuclear-encoded OXPHOS transcript *NDUFA7* (normalized to unlabeled ERCC-0048 spike-in control) from two replicate measurements of total RNA from FASTKD4-deficient cells compared to the control. Steady-state RNA abundance was determined by quantifying unlabeled RNA from cells not treated with 4sU. **C)** Log2 fold change (FASTKD4 KO over NT) in RNA turnover for mtRNAs and nuclear-encoded OXPHOS transcript *NDUFA7* (normalized to unlabeled ERCC-0048 spike-in control) from two replicate measurements of total RNA from FASTKD4-deficient cells compared to the control. RNA turnover was assessed by quantifying the unlabeled RNA remaining after 90 min of labeling with 4sU and dividing by the abundance of unlabeled RNA from cells not treated with 4sU (steady-state abundance). **D)** Poly(A) tail lengths measured by direct RNA sequencing of poly(A)+ mRNA from two replicate measurements of NT control and FASTKD4 KO cells (left). Density plots showing the poly(A) tail length for *ATP8/6* and *ND3* in NT and FASTKD4 KO cells (right). nt, nucleotides. **E)** Comparison of log2 fold change in control-normalized steady-state mtRNA abundance in FASTKD4 KO and LRPPRC KO cells for a subset of mtRNAs colored by OXPHOS complex. Error bars represent standard deviation from two biological replicates. Dashed grey line represents a slope of 1. **F)** NT-normalized transcript abundance (relative to ACTB) for a subset of mt-mRNAs as quantified by RT-qPCR for three replicate measurements from NT, FASTKD4 KO, and LRPPRC KO cells. Error bars represent standard deviation. Brown-Forsythe and Welch ANOVA test followed by Dunnett’s T3 multiple comparisons test, and significant p-values are indicated; *p-value < 0.05, **p-value < 0.01, and ***p-value < 0.001.

**Supplementary Figure 3:**
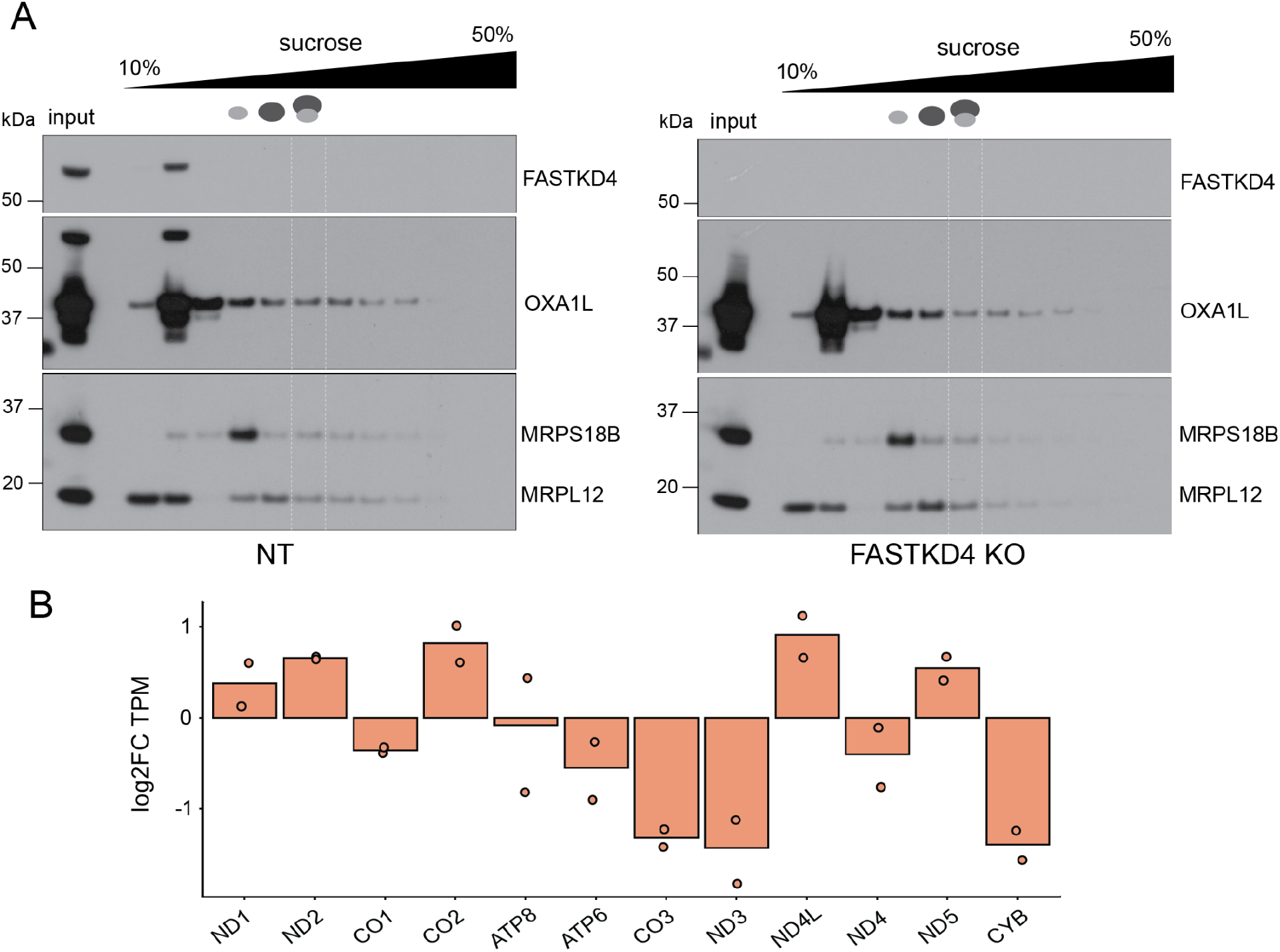
FASTKD4 KO reduces translation of *CYB*. **A)** Isolation of mitomonosome (55S) footprints from mitochondrial polysomes. Lysates were clarified and loaded onto linear 10-50% sucrose gradients. Western blotting against the proteins of the large (MRPL12) and small (MRPS18B) subunits was performed to detect mitoribosomes in each fraction. Representative blots from two replicates are shown. kDa, kilodalton. **B)** Protein synthesis represented by mitoribosome-protected footprints mapping to heavy-strand mt-mRNAs from two replicate measurements of FASTKD4 KO cells compared to the control. TPM was calculated by dividing the RPK, mitoribosome protected fragments normalized by gene length, for each transcript by the sum of RPK values and multiplying by one million. RPF, ribosome-protected footprints;TPM, transcripts per million; RPK, reads per kilobase.

**Supplementary Figure 4:**
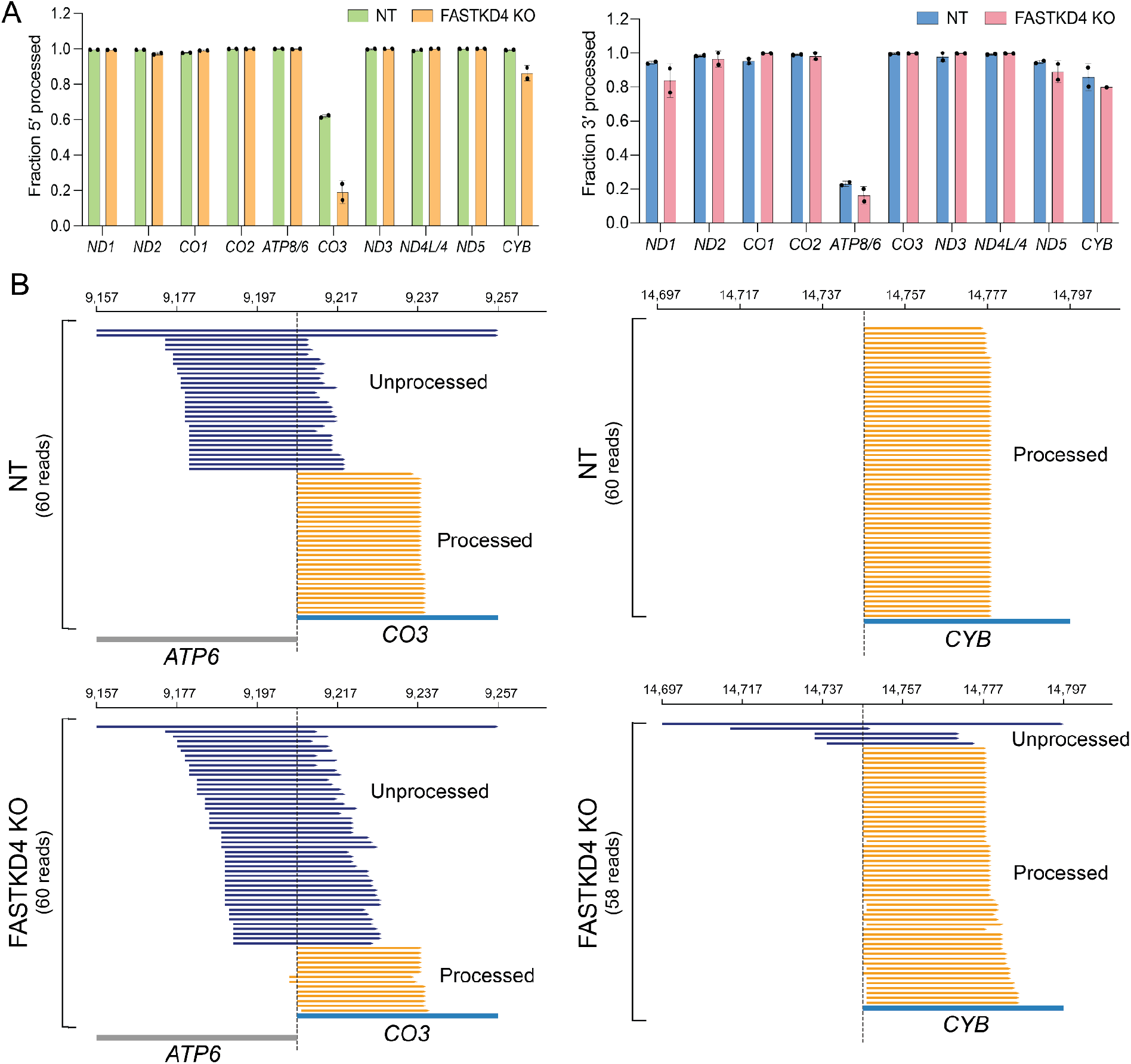
Mitoribosomes predominantely engage 5′ processed transcripts in FASTKD4 KO cells. **A)** Fraction of size-selected RPFs mapping to 5′ processed (left) and 3′ processed (right) heavy-strand transcripts in NT control and FASTKD4 KO cells for two replicate measurements. RPF, ribosome-protected footprint. **B)** Gene track depicting a subsampling of size-selected mitoribosome profiling reads aligned to the *ATP8/6-CO3* (left) and *ND5-CYB* (right) junctions in NT and FASTKD4 KO cells. Dashed line represents the annotated 5′ end of the mature *CO3* or *CYB* transcript.

**Supplementary Figure 5:**
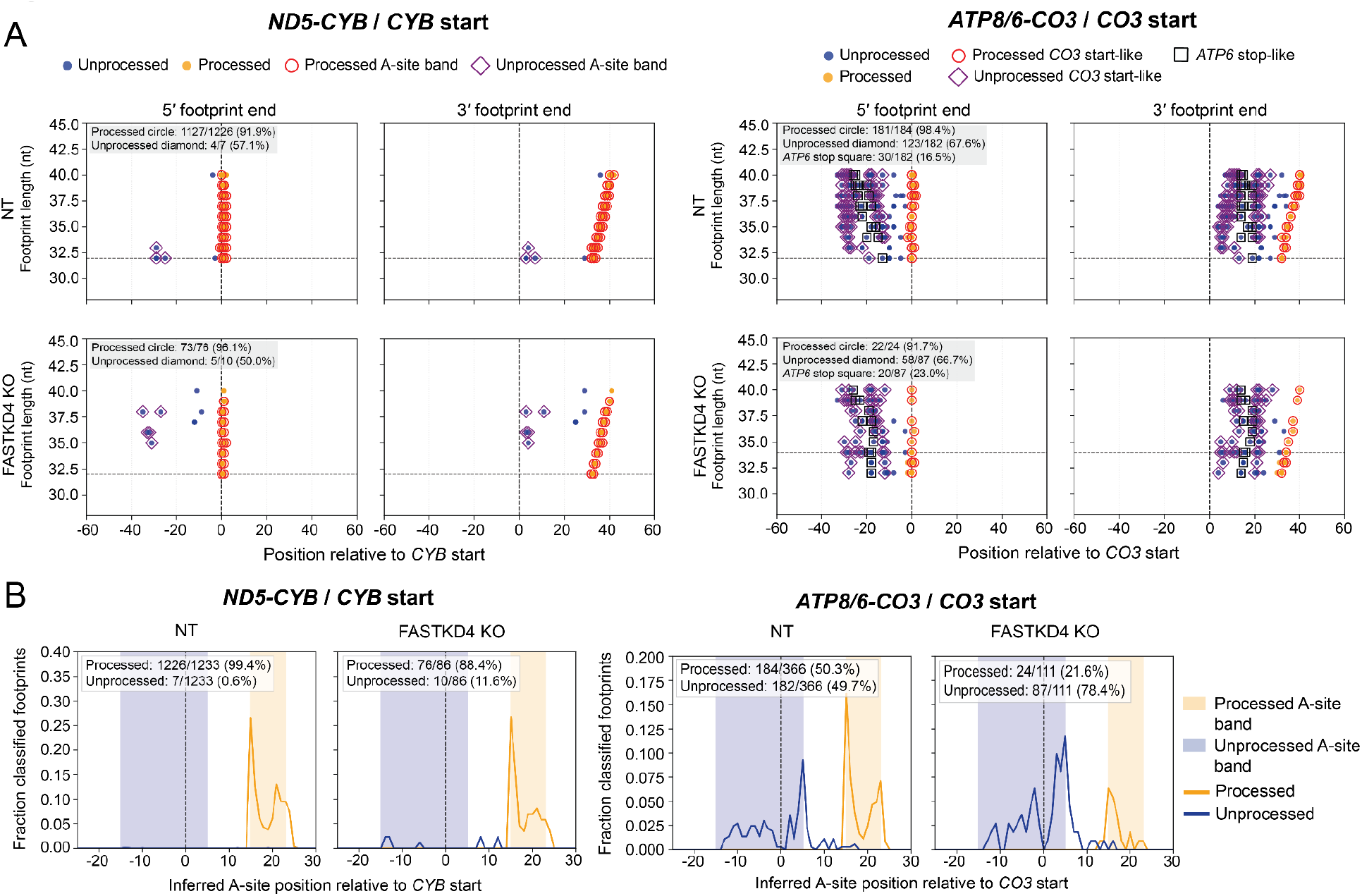
Mitoribosomes selectively engage 5′ processed *CYB*. **A)** Relative positioning of RPF 5′ and 3′ ends, sorted by footprint size, aligned to the *ND5-CYB* (left) and *ATP8/6-CO3* (right) junctions in NT and FASTKD4 KO cells. Dashed line represents the annotated start of the mature *CO3* or *CYB* transcript. Symbols indicate processing status of the read and if the A-site position is start-like. RPF, ribosome protected footprint. **B)** Histogram depicted the fraction of start-like A-site selected RPFs aligned to the *ND5-CYB* (left) and *ATP8/6-CO3* (right) junctions in NT and FASTKD4 KO cells. Dashed line represents the annotated start of the mature *CYB* or *CO3* transcript.

**Supplementary Figure 6:**
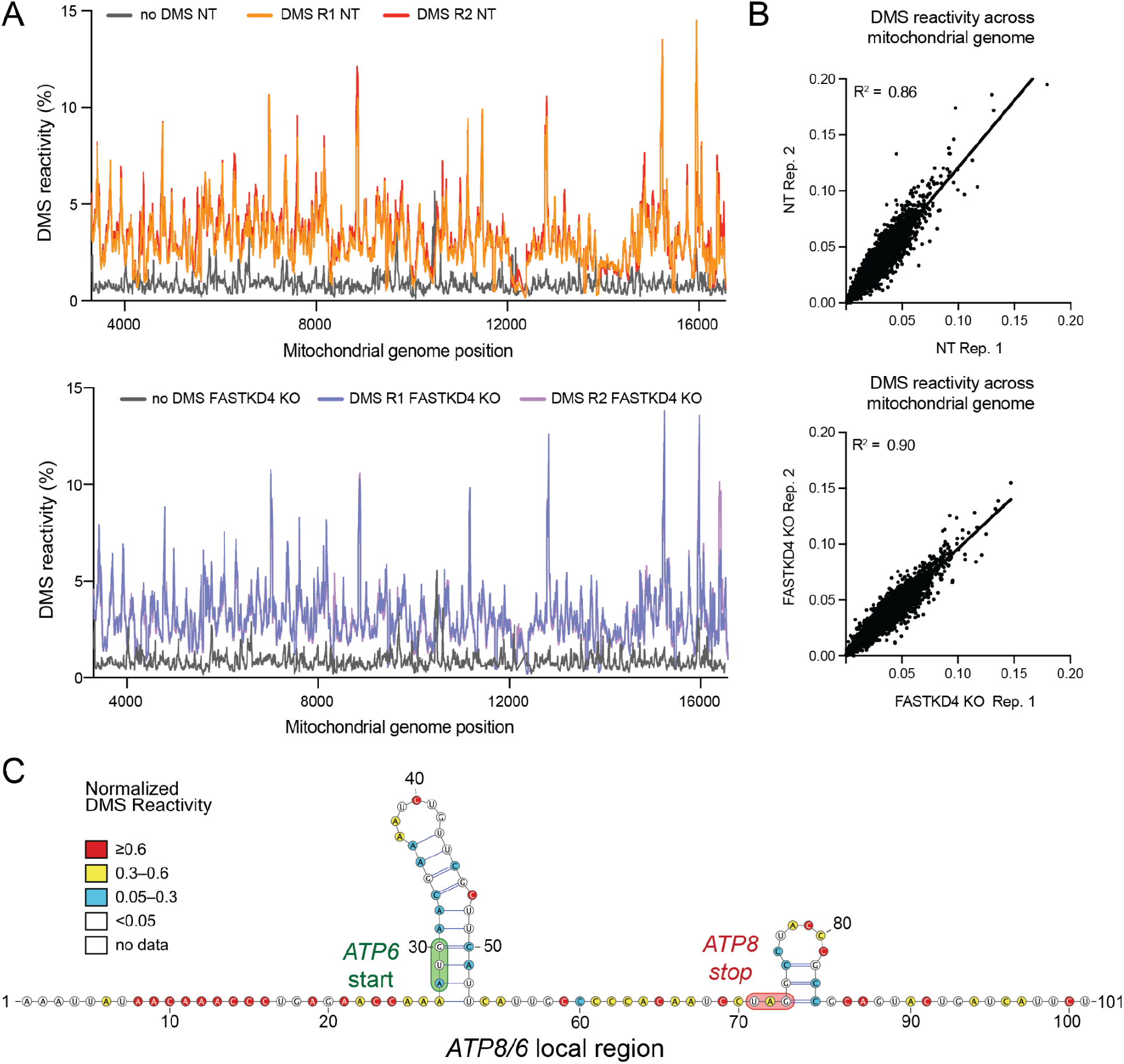
DMS reactivities are well correlated across replicates. **A)** Linear representation of the percent DMS reactivity across the mitochondrial genome for two replicate measurements from DMS-treated NT cells (top) or FASTKD4 KO cells (bottom) compared to no-DMS controls. DMS, dimethyl sulfate. **B)** Interexperimental correlation of percent DMS reactivity across the mitochondrial genome for NT replicates (top) and FASTKD4 KO replicates (bottom). The coefficient of determination (R^2^) value is indicated. **C)** Predicted secondary structure of *ATP8/6* bicistronic transcript in FASTKD4 KO. Structure similar to that observed by Moran et al. (27). Nucleotides are colored by normalized DMS reactivity according to the scale shown. *ATP6* start codon highlighted in green and *ATP8* stop codon highlighted in red.

**Supplementary Figure 7:**
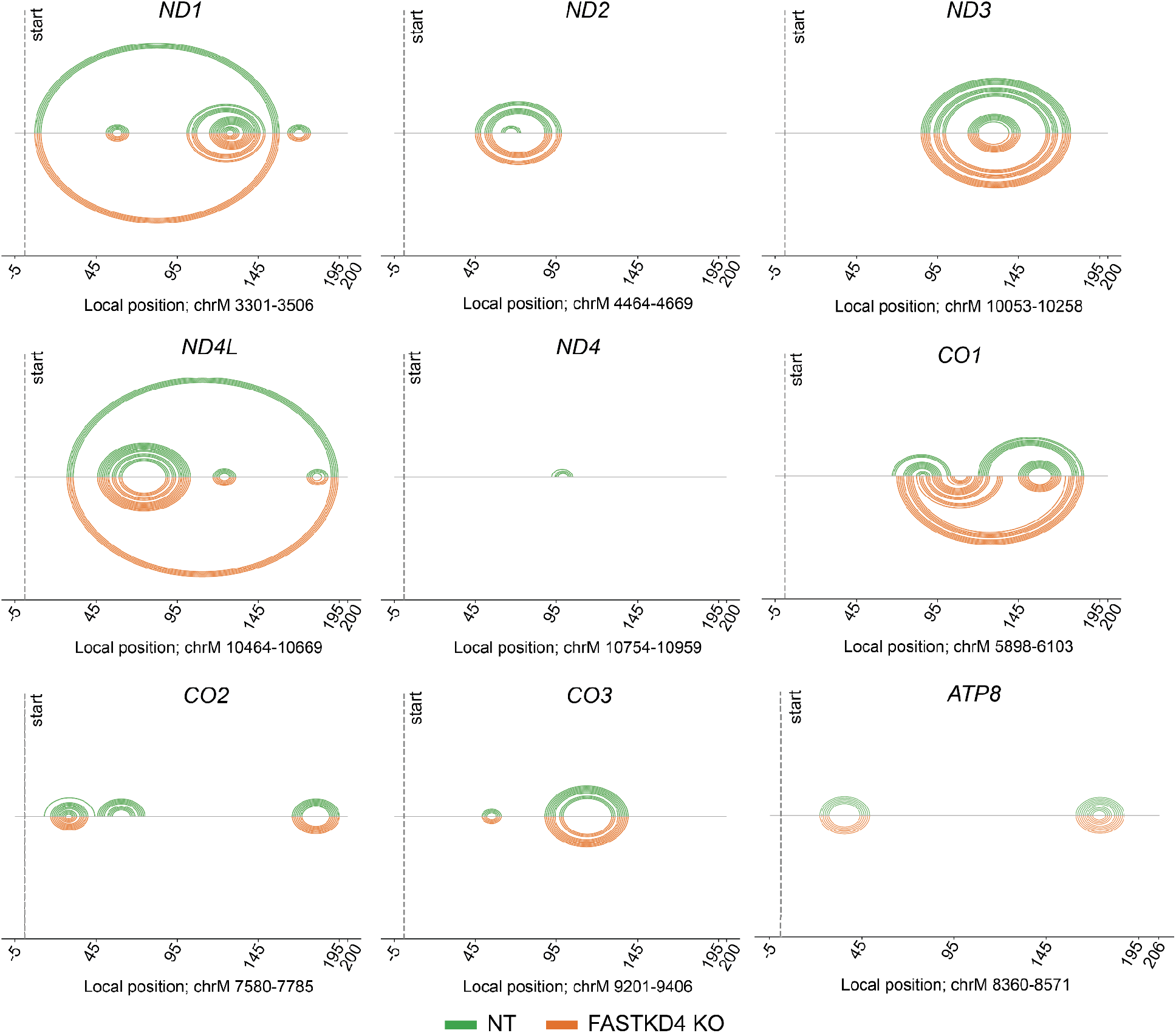
5′-most regions of mt-mRNAs are largely unstructured in control and FASTKD4 KO cells. Linear arc plots depicting predicted changes in RNA secondary structure between NT and FASTKD4 KO within the first 200 nt of heavy-strand mRNAs. nt, nucleotides.

## Notes

### Competing Interest Statement

The authors have declared no competing interest.

