## Supplementary Table 1 for "Mitochondrial RNA processing promotes translation by resolving structured precursor RNAs"

**Supplementary Table S1**

| Reagent | Source | Identifier |
|---|---|---|
| **Antibodies** | | |
| ACTB | Cell Signaling Technology | 3700, RRID:AB_2242334 |
| SDHA | Santa Cruz | sc-166947, RRID:AB_10610526 |
| TBRG4 (FASTKD4) | Sigma | HPA020582, RRID:AB_1857804 |
| MRPS18B | Proteintech | 16139-1-AP, RRID:AB_2146368 |
| MRPL12 | Proteintech | 14795-1-AP, RRID:AB_2250805 |
| MT-ND1 | Abcam | ab181848, RRID:AB_2687504 |
| MT-CO1/COX1 | Abcam | ab14705, RRID:AB_2084810 |
| MT-CO2/COX2 | Proteintech | 55070-1-AP, RRID:AB_10859832 |
| MT-ATP6 | Proteintech | 55313-1-AP, RRID:AB_2881305 |
| MT-CYB/CYTB | Proteintech | 55090-1-AP, RRID:AB_2881266 |
| IRDye 800CW Goat anti-Mouse IgG Secondary Antibody | LICORbio | Cat# 926-32210, RRID:AB_621842 |
| IRDye 800CW Goat anti-Rabbit IgG Secondary Antibody | LICORbio | Cat# 926-32211, RRID:AB_621843 |
| Anti-mouse IgG, HRP-linked Antibody | Cell Signaling Technology | 7076, RRID:AB_330924 |
| Anti-rabbit IgG, HRP-linked Antibody | Cell Signaling Technology | 7074, RRID:AB_2099233 |
| **qRT-PCR probes** | | |
| ACTB | ThermoFisher | Hs03023880_g1 |
| MT-CO1 | ThermoFisher | Hs02596864_g1 |
| MT-CO3 | ThermoFisher | Hs02596866_g1 |
| MT-ND3 | ThermoFisher | Hs02596875_s1 |
| MT-CYB | ThermoFisher | Hs02596867_s1 |
| **Cell lines** | | |
| HEK293T | ATCC | CRL-3216 |
| HeLa | ATCC | CCL-2 |
| HEK293T LRPPRC KO | (Soto et al. 2022) | |

| | | |
|---|---|---|
| HEK293T FASTKD4 KO and control | This paper | |
| HeLa FASTKD4 pooled KO and control | This paper | |
| **Oligonucleotides** | | |
| Forward gRNA for non-targeting AAVS1 control | IDT | GCCAAGGACTCAAACCCAGA |
| Reverse gRNA for non-targeting AAVS1 control | IDT | CCCCGTTCTCCTGTGGATTC |
| Forward gRNA for FASTKD4 KO (sg2) | IDT | CACCGGGTTCATTAGTGGCTCCGAG |
| Reverse gRNA for FASTKD4 KO (sg2) | IDT | AAACCTCGGAGCCACTAATGAACCC |
| Forward gRNA for FASTKD4 KO (sg3) | IDT | CACCGGGAGGTCCGCTGGCGCATG |
| Reverse gRNA for FASTKD4 KO (sg3) | IDT | AAACCATGCGCCAGCGGACCTCCC |
| Forward gRNA for FASTKD4 KO (sg4) | IDT | CACCGGGCCGACTGAGACTTGCC |
| Reverse gRNA for FASTKD4 KO (sg4) | IDT | AAACGGCAAGTCTCAGTCGGCCC |
| **Plasmids** | | |
| pcDNA3.1 | Addgene | V790-20 |
| pcDNA3.1(FASTKD4-FLAG) | This paper | |
| lentiCas9-Blast | Addgene | 52962 |
| lentiGuide-Puro | Addgene | 52963 |
| lentiGuide-Puro (FASTKD4 sg2) | This paper | |
| lentiGuide-Puro (FASTKD4 sg3) | This paper | |
| lentiGuide-Puro (FASTKD4 sg4) | This paper | |
